# Sensing of endogenous nucleic acids by ZBP1 induces keratinocyte necroptosis and skin inflammation

**DOI:** 10.1101/2020.03.25.007096

**Authors:** Michael Devos, Giel Tanghe, Barbara Gilbert, Evelien Dierick, Maud Verheirstraeten, Josephine Nemegeer, Richard de Reuver, Sylvie Lefebvre, Jolien De Munck, Jan Rehwinkel, Peter Vandenabeele, Wim Declercq, Jonathan Maelfait

## Abstract

Aberrant detection of endogenous nucleic acids by the immune system can cause inflammatory disease. The scaffold function of the signalling kinase RIPK1 limits spontaneous activation of the nucleic acid sensor ZBP1. Consequently, loss of RIPK1 in keratinocytes induces ZBP1-dependent necroptosis and skin inflammation. Whether nucleic acid sensing is required to activate ZBP1 in RIPK1 deficient conditions and which immune pathways are associated with skin disease remained open questions. Using knock-in mice with disrupted ZBP1 nucleic acid binding activity, we report that sensing of endogenous nucleic acids by ZBP1 is critical in driving skin pathology characterised by antiviral and IL-17 immune responses. Inducing ZBP1 expression by interferons triggers necroptosis in RIPK1-deficient keratinocytes and epidermis-specific deletion of MLKL prevents disease, demonstrating that cell-intrinsic events cause inflammation. These findings indicate that dysregulated sensing of endogenous nucleic acid by ZBP1 can drive inflammation and may contribute to the pathogenesis of IL-17-driven inflammatory skin conditions such as psoriasis.

**Summary:** Devos, Tanghe *et al*. find that the recognition of endogenous nucleic acids by the nucleic acid sensor ZBP1 causes necroptosis of RIPK1-deficient keratinocytes. This process drives the development of an inflammatory skin disease characterised by an IL-17 immune response.

## Introduction

Erroneous detection of endogenous nucleic acids by pattern recognition receptors (PRRs) of the innate immune system can cause the development of inflammatory diseases. This phenomenon is best described in the context of a group of human autoinflammatory and autoimmune disorders, termed type I interferonopathies (Crowl et al., 2017; Lee-Kirsch, 2017; Rodero and Crow, 2016). Gain-of-function mutations in genes encoding components of the nucleic acid sensing pathways, including *TMEM173* (STING) and *IFIH1* (MDA5), result in enhanced detection of endogenous nucleic acids (Jeremiah et al., 2014; Liu et al., 2014; Rice et al., 2014). Conversely, loss-of-function of genes controlling nucleic acid metabolism such as *TREX1, SAMHD1* and *ADAR* result in the reduced clearance of endogenous nucleic acids and triggers spontaneous production of type I interferons (IFNs), which initiate and sustain inflammatory disease development (Crow et al., 2006; Rice et al., 2009; Rice et al., 2012).

The nucleic acid receptor Z-DNA binding protein 1 (ZBP1) restricts RNA and DNA virus infection by mechanisms including the induction of regulated necroptotic cell death (Kuriakose and Kanneganti, 2018). Although regulated cell death is a vital antiviral defence strategy, excessive cell death is thought to contribute to the pathogenesis of inflammatory diseases (Newton and Manning, 2016; Pasparakis and Vandenabeele, 2015; Weinlich et al., 2017). Execution of necroptosis results in cell membrane rupture and the release of intracellular components including damage-associated molecular patterns (DAMPs), which activate innate immune receptors thereby initiating a detrimental inflammatory response. The release of antigens by dying cells may promote adaptive immune responses further driving autoimmune pathology and establishing chronic disease (Galluzzi et al., 2017).

In mouse cells, the induction of necroptosis following ZBP1 engagement occurs via the direct recruitment of the serine/threonine protein kinase RIPK3 through RIP homotypic interaction motifs (RHIMs), which are present in both RIPK3 and ZBP1 (Upton et al., 2012). RIPK3 then phosphorylates the pseudokinase MLKL on serine 345, leading to its oligomerisation and translocation to the plasma membrane, where it damages the integrity of the cell membrane (Wallach et al., 2016). In contrast to necroptosis triggered by tumor necrosis factor (TNF), necroptosis downstream of ZBP1 does not require the kinase activity of the RHIM-containing signalling kinase RIPK1 (Upton et al., 2012). In keratinocytes, ablation of RIPK1 or disruptive mutation of the RHIM of RIPK1 induces spontaneous ZBP1-dependent necroptosis and triggers skin inflammation, indicating that RIPK1 acts as a molecular scaffold to inhibit ZBP1-RIPK3-MLKL signalling in a RHIM-dependent way (Dannappel et al., 2014; Lin et al., 2016; Newton et al., 2016).

ZBP1 contains two tandem N-terminal Z-form nucleic acid binding (Zα)-domains, which specifically interact with double stranded (ds) nucleic acid helices in the Z-conformation, including Z-RNA and Z-DNA (Herbert, 2019). Others and our group have shown that engagement of ZBP1 upon virus infection crucially depended on nucleic acid sensing by intact Zα-domains (Maelfait et al., 2017; Sridharan et al., 2017; Thapa et al., 2016). The identity of viral ZBP1 agonists remains uncertain, and viral ribonucleoprotein complexes, RNA genomes or viral transcripts have been suggested to interact with the Zα-domains of ZBP1 (Guo et al., 2018; Kesavardhana et al., 2017; Maelfait et al., 2017; Sridharan et al., 2017; Thapa et al., 2016). The importance of Z-form nucleic acid interaction with ZBP1 during viral infection raises the question whether endogenous nucleic acids can also stimulate ZBP1 when certain checkpoints such as RIPK1 are compromised and whether aberrant nucleic acid sensing by ZBP1 could provoke the development of inflammatory disease.

To address this question, we crossed epidermis-specific RIPK1-deficient mice, which developed inflammatory skin disease, to *Zbp1* knock-in animals with mutated Zα-domains, rendering ZBP1 unable to interact with Z-form nucleic acids (Maelfait et al., 2017). Using this genetic approach, we show that skin pathology critically depends on nucleic acid sensing by ZBP1. The epidermis of keratinocyte-specific RIPK1-deficient mice displayed enhanced expression of antiviral genes, including ZBP1, and is characterised by infiltration of IL-17-producing CD4 T cells and innate lymphoid cells (ILCs). Finally, *in vitro* type I (IFNβ) or type II IFN (IFNγ) treatment of RIPK1-deficient keratinocytes induced necroptosis, while RIPK1-deficient cells expressing Zα-domain mutant ZBP1 were protected. Together, these data suggest that endogenous nucleic acid sensing by ZBP1 triggers cell-intrinsic necroptosis of RIPK1-deficient keratinocytes, which initiates an inflammatory signalling cascade causing inflammatory skin pathology.

## Results and Discussion

### Skin pathology of *Ripk1*^EKO^ mice is dependent on nucleic acid sensing by ZBP1

Mice with epidermis-specific deletion of RIPK1 develop progressive ZBP1-dependent skin inflammation (Dannappel et al., 2014; Lin et al., 2016). To determine if nucleic acid sensing by ZBP1 is required for skin pathology, we crossed *Ripk1*^FL/FL^ K5Cre^Tg/+^ animals (*Ripk1*^EKO^) (Ramirez et al., 2004; Takahashi et al., 2014) to mice carrying a *Zbp1* allele (*Zbp1*^Zα1α2^) mutated in the two Zα-domain coding regions, resulting in the expression of a ZBP1 protein which is unable to interact with its nucleic acid agonist. We have previously shown that *Zbp1*^Zα1α2/Zα1α2^ mice are susceptible to a strain of murine cytomegalovirus that does not block necroptosis downstream of ZBP1 (Maelfait et al., 2017). *Ripk1*^EKO^ mice developed skin pathology with macroscopically visible lesions starting 1 week after birth (Fig. 1A and Fig. S1A). In contrast, *Ripk1*^EKO^ *Zbp1*^Zα1α2/Zα1α2^ animals remained lesion-free until at least 12 weeks after birth (Fig. 1A), whereupon more than half of the mice displayed smaller and more focal lesions compared to *Ripk1*^EKO^ littermates and these lesions did not develop further until 1 year of age. One intact allele of *Zbp1* was sufficient to initiate disease development, albeit with delayed kinetics (Fig. 1A). ZBP1 protein levels were not affected by Zα-domain mutation or the absence of RIPK1 in primary keratinocytes treated with IFNβ to induce the expression of ZBP1 (Fig. S1B). Importantly, the interaction between ZBP1 and RIPK3 was not disrupted due to Zα-domain mutation as RIPK3 co-immunoprecipitated equally efficient with wild type and mutant ZBP1 (Fig. S1C). Absence of inflammation was confirmed by histological examination of skin of 4 to 5 week old *Ripk1*^EKO^ *Zbp1*^Zα1α2/Zα1α2^ mice, as shown by normal epidermal thickness (Fig 1B and 1C) and a normal expression pattern of the epidermal differentiation markers keratin-1, -5 and -6, compared to *Ripk1*^EKO^ mice (Fig 1B). As previously described, *Ripk1*^EKO^ mice crossed to full-body *Zbp1* knock-out animals displayed a similarly rescued phenotype as the *Ripk1*^EKO^ *Zbp1*^Zα1α2/Zα1α2^ line (Fig S1D) (Lin et al., 2016). Deletion of *Ripk3* or reducing its gene dosage by half was sufficient to establish substantial protection, confirming that activation of RIPK3 led to skin pathology (Fig S1E) (Dannappel et al., 2014). Loss of RIPK3 offered better protection against lesion formation than ZBP1 Zα-domain mutation or deficiency, indicating that other RHIM-containing signalling complexes, most likely mediated by TRIF, contribute to disease progression (Dannappel et al., 2014). We conclude that intact Zα-domains are crucial for ZBP1-induced lesion formation in *Ripk1*^EKO^ mice, indicating that nucleic acid sensing by ZBP1 drives keratinocyte necroptosis in absence of RIPK1.

**Figure 1.**
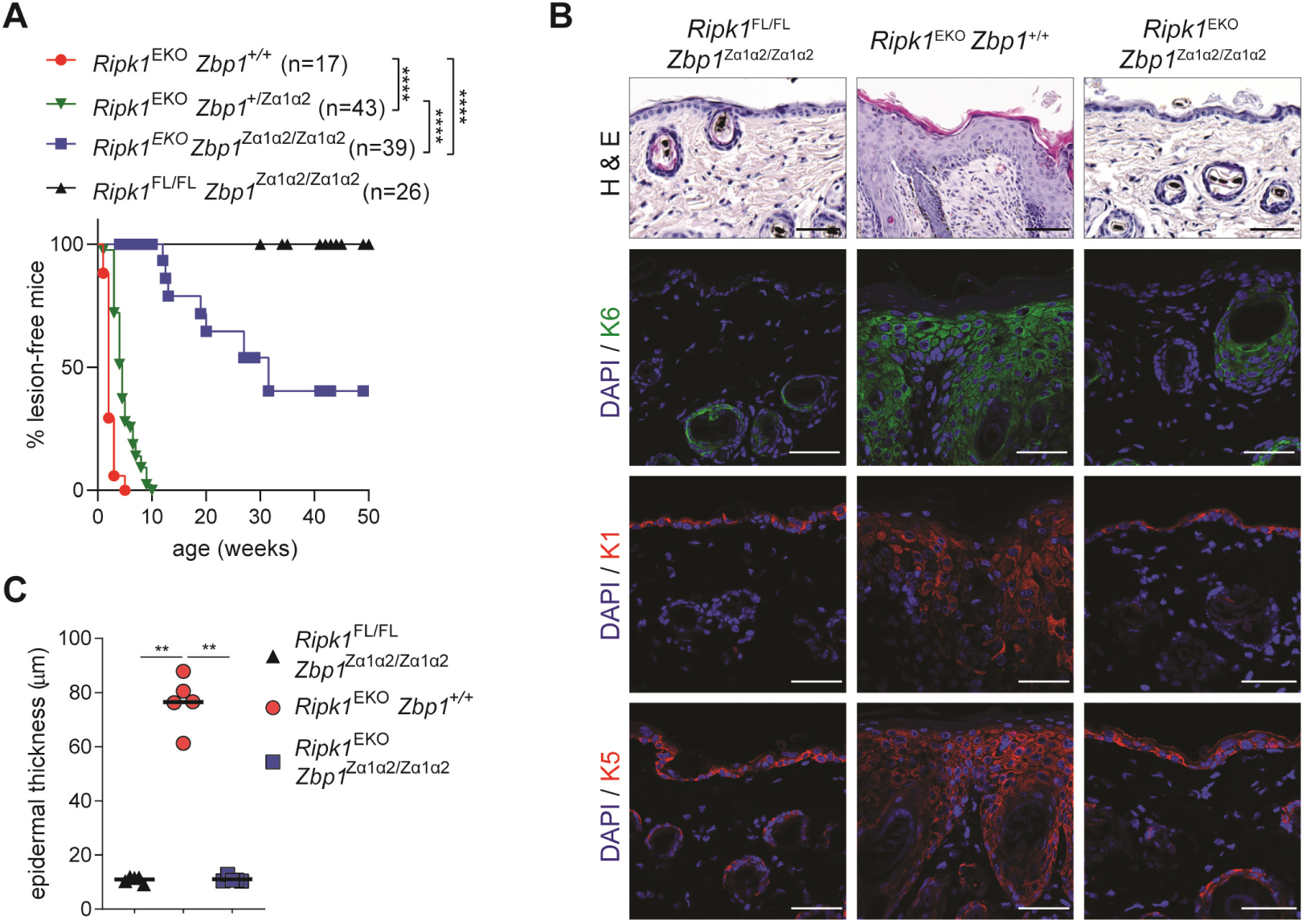
Skin pathology of *Ripk1*^EKO^ mice is dependent on nucleic acid sensing by ZBP1. **(A)** Kaplan-Meier plot of macroscopically visible lesion appearance of epidermis-specific RIPK1 deficient mice (*Ripk1*^EKO^), *Ripk1*^EKO^ mice carrying heterozygous (*Zbp1*^+/Zα1α2^) or homozygous (*Zbp1*^Zα1α2/Zα1α2^) *Zbp1* Zα-domain mutant alleles. Littermates which did not express the keratinocyte-specific K5-Cre transgene (*Ripk1*^FL/FL^ *Zbp1*^Zα1α2/Zα1α2^) were used as controls. Pictures of lesions are shown in supplemental figure S1A. ****p < 0.0001 by Log-Rank test. **(B)** Back skin sections of 4 to 5 week old mice with indicated genotypes were stained with H&E or immunostained for keratin 6 (K6), keratin 1 (K1) or keratin 5 (K5) antibodies and DAPI. Scale bars represent 50 μm. At least 4 mice per genotype were analysed. **(C)** Quantification of epidermal thickness on H&E-stained sections shown in **(B)**. The line represents the mean and dots represent individual mice. **p < 0.01 by Mann-Whitney U-test. Data are representative of at least two independent experiments.

### ZBP1 activation drives an IL-17 immune response in *Ripk1*^EKO^ mice

Next, we determined the presence of immune cells in inflammatory lesions of *Ripk1*^EKO^ mice by flow cytometry (Fig. S2A). The total number of CD45^+^ leukocytes was significantly increased in *Ripk1*^EKO^ epidermis and this was dependent on nucleic acid sensing by ZBP1 (Fig. 2A and S2A). In healthy mouse skin γδ-TCR^+^ dendritic epidermal T cells (DETCs) constitute the majority of leukocytes (Nielsen et al., 2017), however, their numbers did not significantly increase in RIPK1-deficient skin (Fig. 2A and S2A). In contrast, RIPK1-deficient epidermis displayed a marked infiltration of αβ-TCR^+^ CD4 T cells, which reached similar levels as DETCs. The accumulation of CD4 T cells depended on intact nucleic acid sensing by ZBP1 as these cells were reduced to wild type levels in the epidermis of *Ripk1*^EKO^ *Zbp1*^Zα1α2/Zα1α2^ mice (Fig. 2A and S2A). Further quantification showed a ZBP1-dependent influx of CD8 T cells, NKT cells, neutrophils and CD64^+^ myeloid cells (Fig. 2A and S2A). *In vitro* activation assays on cells isolated from the epidermis of *Ripk1*^EKO^ mice revealed that the majority of the CD4 T cells produced the cytokine IL-17A, but not IFNγ, identifying these cells as Th17 cells (Fig 2B and 2C). A large fraction of the CD45^+^ leukocytes stained negative for many of the cell surface markers used in our staining panel, however, they produced large amounts of IL-17A and equalled Th17 cells in total cell numbers (Fig. 2B and 2C). We concluded that these cells most likely represent type 3 innate lymphoid cells (ILC3s). At 5 weeks of age, the infiltration of IL-17A-producing CD4 T cells and ILC3s was not observed in *Ripk1*^EKO^ *Zbp1*^Zα1α2/Zα1α2^ double transgenic mice. However, the epidermis of 10 to 12 week old *Ripk1*^EKO^ *Zbp1*^Zα1α2/Zα1α2^ animals contained increased numbers of Th17 and ILC3s (Fig. 2C), which coincided with the development of mild skin pathology and correlated with the incomplete phenotypic rescue of these mice (see Fig. 1A). In addition, we detected ZBP1-dependent increased messenger RNA expression of the IL-17A regulatory cytokine IL-23 in RIPK1-deficient epidermis (Fig. 2D). Together, these results indicate that in the absence of RIPK1, the detection of nucleic acids by ZBP1 triggers keratinocyte necroptosis, which induces an IL-17-mediated inflammatory response.

**Figure 2.**
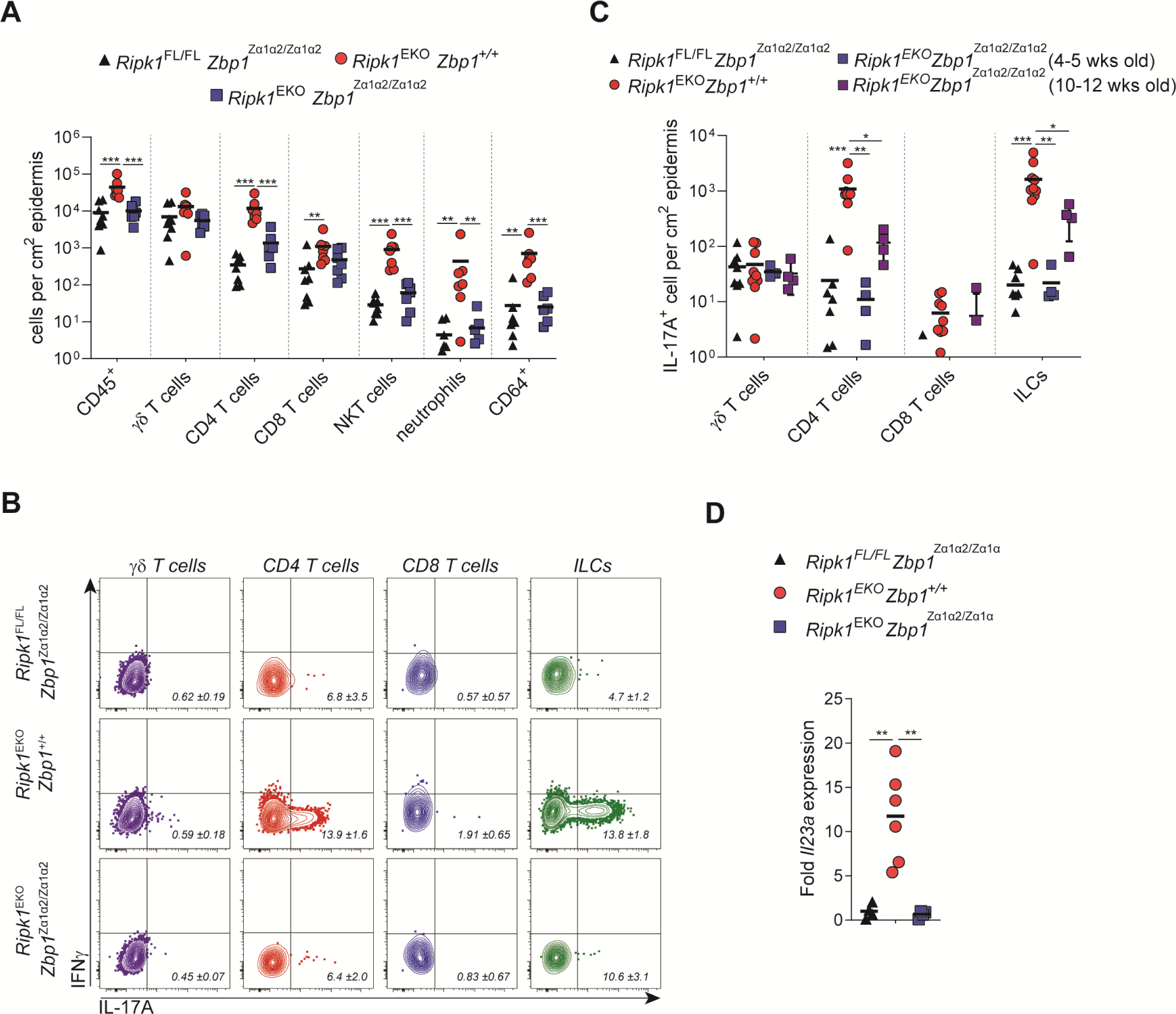
ZBP1 activation drives antiviral an IL-17 immune response in *Ripk1*^EKO^ mice. **(A**) Flow cytometry analysis of leukocyte (CD45^+^) composition of epidermis of mice of the indicated genotypes. The number of immune cells per cm^2^ of epidermis are plotted. Each dot represents an individual mouse and the line represents the mean. The gating strategy is outlined in supplemental figure S2B. **(B and C)** Epidermal cells isolated from 4 to 5 week *Ripk1*^FL/FL^ *Zbp1*^Zα1α2/Zα1α2^, *Ripk1*^EKO^ *Zbp1*^+/+^ or *Ripk1*^EKO^ *Zbp1*^Zα1α2/Zα1α2^ or 10 to 12 week old *Ripk1*^EKO^ *Zbp1*^Zα1α2/Zα1α2^ mice were activated with PMA and ionomycin and stained for intracellular expression of IFNγ and IL-17A. In panel **(B)** representative flow cytometry plots are shown for each genotype. Panel **(C)** represents the number of IL-17A positive immune cells per cm^2^ of epidermis. Each dot represents an individual mouse and the line represents the mean. **(D)** RT-qPCR analysis of *Il23a* in whole back skin of 4 to 5 week old mice of the indicated genotypes. Data in (A), (B), (C) and (D) are representative of 2 independent experiments. **p < 0.01, ***p < 0.001 by Mann-Whitney U-test.

### Induction of MLKL-dependent necroptosis by ZBP1 in keratinocytes drives skin inflammation in *Ripk1*^EKO^ mice

ZBP1 expression is inducible by type I and type II IFNs and its expression is increased in RIPK1-deficient epidermis (Lin et al., 2016). We therefore examined whether skin lesions of *Ripk1*^EKO^ mice showed a general antiviral gene signature. Indeed, messenger RNA expression of a panel of interferon stimulated genes (ISGs; *Zbp1, Ifi44, Isg15* and *Ifit1*) and type II IFN (*Ifng*) was enhanced in 5 week old *Ripk1*^EKO^ mice and this was fully restored in a ZBP1 Zα-domain mutant background (Fig. 3A and Fig S2B). In addition, increased expression of inflammatory cytokines (*Il6, Il1b, Il1a, Il33*), the chemokine *Ccl20* and antimicrobial peptide *S100a8* required intact Zα-domains of ZBP1 (Fig. 3A and Fig S2B). To determine in which cell types ZBP1 was expressed in inflamed *Ripk1*^EKO^ skin, we performed immunostaining of ZBP1. In normal skin, ZBP1 expression is restricted to a few dermal cells, probably myeloid cells such as macrophages, which express high basal levels of ZBP1 (Lin et al., 2016; Newton et al., 2016). In contrast, inflamed epidermis from *Ripk1*^EKO^ mice displayed strong cytosolic staining for ZBP1 in keratinocytes (Fig. 3B), consistent with the enhanced ZBP1 messenger RNA expression in these lesions (Fig. 3A). The high expression of ZBP1 in RIPK1-deficient keratinocytes could induce MLKL-dependent necroptosis in these cells to drive skin inflammation in *Ripk1*^EKO^ mice. To test this hypothesis, we crossed *Ripk1*^EKO^ mice to *Mlkl*^FL/FL^ animals (Murphy et al., 2013), generating mice lacking both RIPK1 and MLKL in keratinocytes. Similar to complete ZBP1 deficiency, ablating MLKL specifically in keratinocytes profoundly attenuated the development of lesion formation (Fig. 3C). In addition to inducing necroptosis, ZBP1 engagement has been reported to activate the NLRP3 inflammasome leading to caspase-1-dependent pyroptosis and IL-1β release (Kuriakose et al., 2016), which may further contribute to skin inflammation. However, *Ripk1*^EKO^ *Casp1/11*^−/−^ double transgenic mice were indistinguishable from *Ripk1*^EKO^ littermates in terms of skin lesion development thereby excluding a role for pyroptosis in skin pathology (Fig. 3D). Together, these results show that sensing of nucleic acids by ZBP1 in keratinocytes of *Ripk1*^EKO^ mice causes MLKL-dependent necroptosis, which initiates skin inflammation.

**Figure 3.**
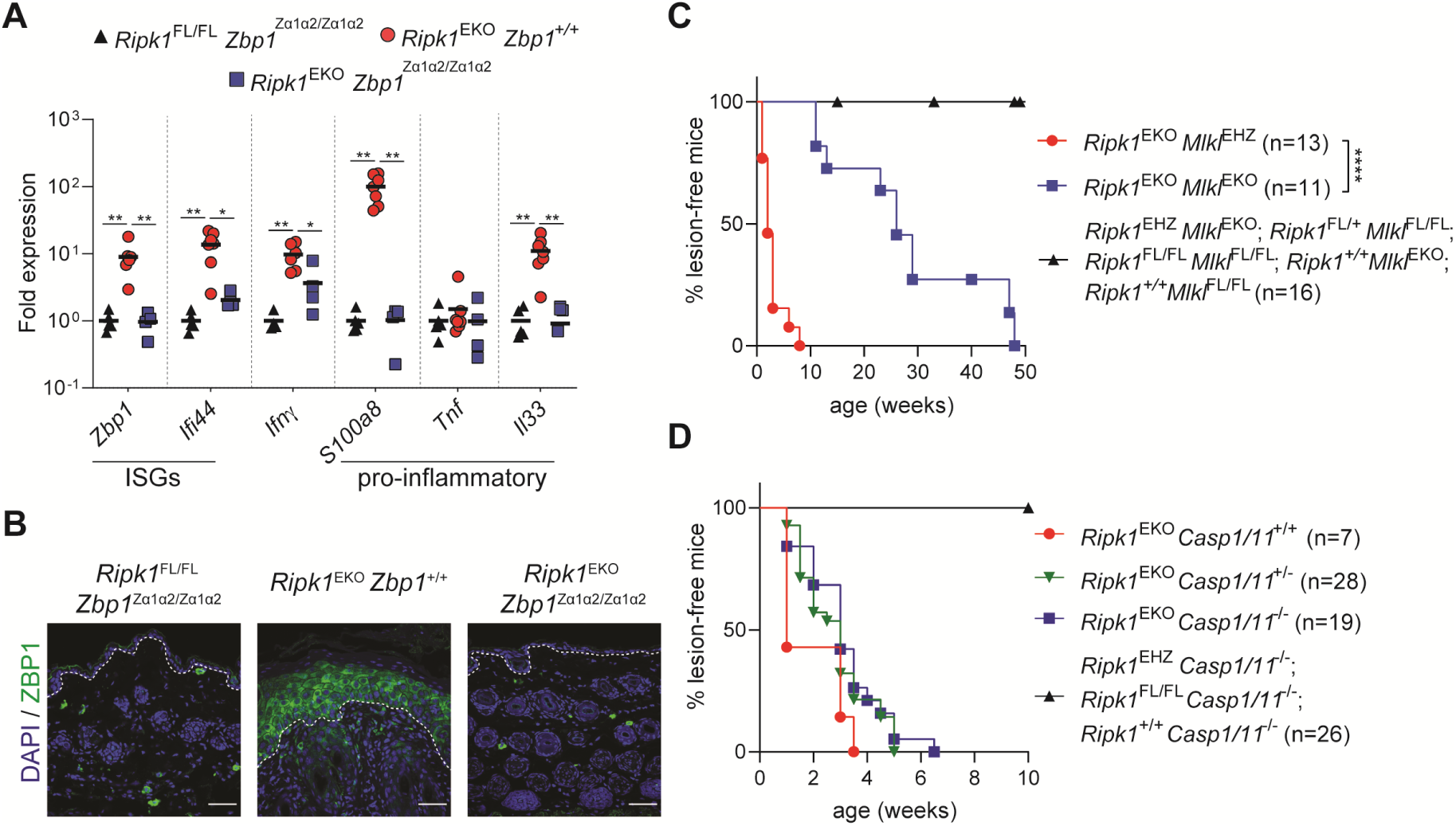
Induction of MLKL-dependent necroptosis by ZBP1 in keratinocytes drives skin inflammation in *Ripk1*^EKO^ mice. **(A)** RT-qPCR analysis of indicated interferon-stimulated genes (ISGs), IFNγ and pro-inflammatory genes in whole back skin of 4 to 5 week old mice of the indicated genotypes. The line represents the mean and dots represent individual mice. *p < 0.05 and **p < 0.01 by Mann-Whitney U-test. **(B)** Immunostaining with anti-ZBP1 on back skin sections from 5 to 7 week old *Ripk1*^FL/FL^ *Zbp1*^Zα1α2/Zα1α2^, *Ripk1*^EKO^ *Zbp1*^+/+^ or *Ripk1*^EKO^ *Zbp1*^Zα1α2/Zα1α2^ mice. Scale bars represent 50 μm. The dotted line indicates the border between the epidermis and dermis. At least 4 mice per genotype were analysed. **(C)** Kaplan-Meier plot of macroscopically visible lesion appearance of epidermis-specific RIPK1 and MLKL double deficient mice (*Ripk1*^EKO^ *Mlkl*^EKO^). *Ripk1*^EKO^ *Mlkl*^EHZ^ mice, heterozygously expressing a functional *Mlkl* allele in the epidermis in a *Ripk1*^EKO^ background, developed lesions at the same rate as *Ripk1*^EKO^ mice, as shown in figure 1A. Littermate offspring of the indicated genotypes containing one or two functional *Ripk1* and/or *Mlkl* alleles did not develop lesions and are shown as controls. ****p < 0.0001 by Log-Rank test. **(D)** Kaplan-Meier plot of macroscopically visible lesion appearance of epidermis-specific RIPK1 and full-body Caspase1/11 double deficient mice (*Ripk1*^EKO^ *Casp1/11*^−/−^). Caspase-1/11-sufficient *Ripk1*^EKO^ *Casp1/11*^+/+^ or *Ripk1*^EKO^ *Casp1/11*^+/-^ mice developed lesions at the same rate as *Ripk1*^EKO^ *Casp1/11*^−/−^. Littermate offspring of the indicated genotypes expressing one or two functional *Ripk1* alleles did not develop lesions and are shown as controls. Data shown in (A) and (B) are representative of at least 2 independent experiments.

### ZBP1 activation by endogenous nucleic acids induces necroptosis in RIPK1-deficient keratinocytes

To establish if ZBP1 directly induced cell death of RIPK1-deficient keratinocytes, we treated primary RIPK1-deficient keratinocytes with type I (IFNβ) or type II IFNs (IFNγ), which strongly induced ZBP1 protein levels (Fig. 4A and Fig. S1B), and monitored cell death by measuring uptake of a cell impermeable dye over a 48-hour time course. At 12 hours post treatment, RIPK1-deficient cells (*Ripk1*^EKO^ *Zbp1*^+/+^) started to die, whereas RIPK1-sufficient *Ripk1*^FL/FL^ *Zbp1*^+/+^ or *Ripk1*^FL/FL^ *Zbp1*^Zα1α2/Zα1α2^ keratinocytes were unaffected (Fig. 4C). Immunoblotting revealed that IFNγ or IFNβ treatment induced the phosphorylation of MLKL and RIPK3 only in cells that were lacking RIPK1 and expressing wild type ZBP1 (Fig. 4A). In contrast, keratinocytes derived from *Ripk1*^EKO^ *Zbp1*^Zα1α2/Zα1α2^ mice, were fully resistant to IFNγ- or IFNβ-induced cell death and did not show phosphorylation of MLKL and RIPK3 (Fig. 4A and 4C). Lentiviral transduction of wild type but not Zα-domain mutant ZBP1 (ZBP1^Zα1α2mut^) caused increased cell death of RIPK1-deficient keratinocytes starting at 24 hours post transduction, compared to the empty vector control (Fig. S3A and S3B). These data suggest that endogenous nucleic acids activate ZBP1 in the absence of RIPK1. TNF stimulation did not induce cell death or MLKL and RIPK3 phosphorylation in RIPK1-deficient keratinocytes and treatment with poly(I:C), an agonist for TLR3 which signals via TRIF, only modestly sensitised cells to necroptosis and independently of nucleic acid sensing by ZBP1 (Fig. 4B and 4C). These results suggest that RIPK1 deficiency greatly sensitises keratinocytes to ZBP1-induced necroptosis, but not downstream of other RHIM-containing signalling complexes formed after TNFRI stimulation and only modestly upon TLR3 engagement. The sensitisation of RIPK1-deficient keratinocytes to TLR3/TRIF-mediated necroptosis may explain the mild amelioration of skin inflammation in *Ripk1*^EKO^ mice by epidermis-specific deletion of TRIF (Dannappel et al., 2014). As a control, treatment with a combination of TNF and the pan-caspase inhibitor zVAD, which induces TNFRI-dependent necroptosis, only induced MLKL and RIPK3 phosphorylation and cell death in RIPK1-sufficient cells (Fig. 4D and Fig. S3C). Loss of RIPK1 sensitised keratinocytes to caspase-8-dependent apoptosis induced by treating cells with TNF and the protein synthesis inhibitor cycloheximide (CHX) (Fig. 4D and Fig. S3C), which is in agreement with previous reports (Gentle et al., 2011; Kelliher et al., 1998). Spontaneous ZBP1 activation in RIPK1-deficient keratinocytes did not affect IFNγ-induced gene expression as ISG messenger RNA expression or CXCL10 protein production did not differ between the genotypes (Fig. S3D and S3E).

**Figure 4.**
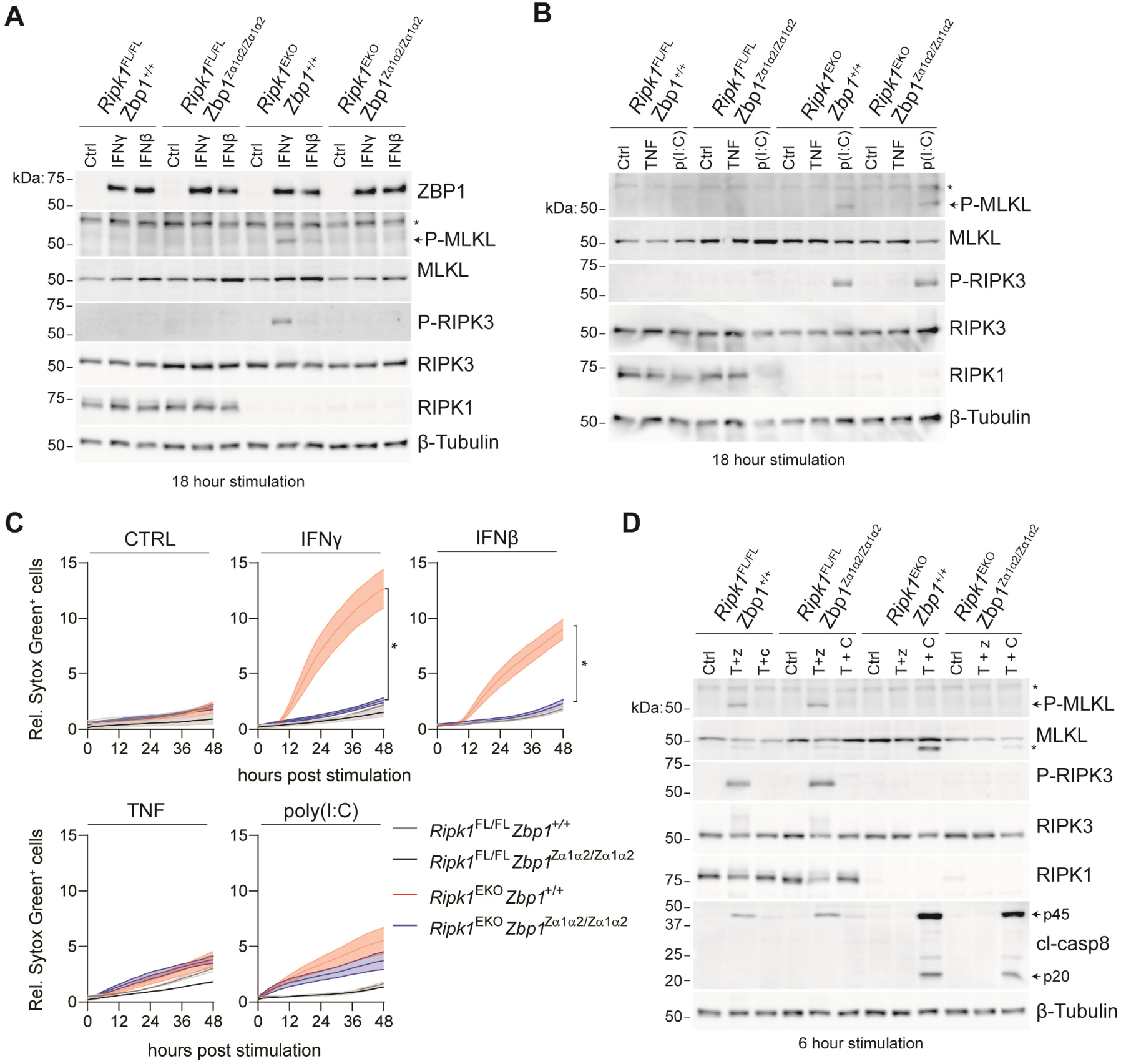
ZBP1 activation by endogenous nucleic acids induces necroptosis in RIPK1-deficient keratinocytes. **(A), (B) and (D)** Primary keratinocytes isolated from *Ripk1*^FL/FL^ *Zbp1*^+/+^, *Ripk1*^FL/FL^ *Zbp1*^Zα1α2/Zα1α2^, *Ripk1*^EKO^ *Zbp1*^+/+^ or *Ripk1*^EKO^ *Zbp1*^Zα1α2/Zα1α2^ mice were stimulated for 18 hours with 200 U/ml IFNγ, 200 U/ml IFNβ, 30 ng/ml TNF or 25 µg/ml poly(I:C) [p(I:C)] or for 6 hours with 30 ng/ml TNF with 20 µM zVAD-fmk (T + z) or 30 ng/ml TNF with 5 µg/ml cycloheximide (T + C). Protein expression was analysed by Western blotting. The asterisk (*) on the phospho(P)-MLKL blot indicates a non-specific signal. The cleaved caspase-8 (cl-casp8) forms p45 and p20 are indicated with arrows. **(C)** Analysis of cell death upon stimulation with 200 U/ml IFNγ, 200 U/ml IFNβ, 30 ng/ml TNF or 25 µg/ml poly(I:C) by measuring Sytox Green uptake every 2 hour of primary keratinocytes isolated from mice of the indicated genotype. Solid lines represent the mean of cell death curves of 2 to 4 primary keratinocyte cultures isolated from different mice. Shaded areas indicate SEM. *p < 0.05 by 2way ANOVA on data from *Ripk1*^EKO^ *Zbp1*^+/+^ (n=4) and *Ripk1*^EKO^ *Zbp1*^Zα1α2/Zα1α2^ (n=4) ultures. Data shown in (A), (B), (C) and (D) are representative of at least 2 independent experiments.

In summary, we report that ZBP1 induces a cell-intrinsic ZBP1-RIPK3-MLKL pro-necroptotic signalling cascade in RIPK1-deficient keratinocytes resulting in inflammatory skin disease. Disease development in *Ripk1*^EKO^ mice depended on the nucleic acid sensing capacity of ZBP1, supporting the idea that endogenous nucleic acids function as the upstream trigger for the execution of necroptosis. The molecular identity of the nucleic acid agonist of ZBP1 under these conditions remains unknown. Two lines of evidence favour the hypothesis that cellular nucleic acids, and not those of viral or bacterial origin, activate ZBP1. Firstly, cultured primary RIPK1 keratinocytes, grown in sterile conditions, succumb to IFN-induced necroptosis, which crucially depended on the nucleic acid sensing capability of ZBP1. This is in agreement with earlier studies reporting on the toxicity of IFNs for RIPK1-deficient fibroblasts (Dillon et al., 2014; Kaiser et al., 2014), and which was recently shown to depend on ZBP1 (Ingram et al., 2019; Yang et al., 2019). Secondly, embryos expressing a mutant RIPK1 RHIM develop ZBP1-RIPK3-MLKL driven inflammatory epidermal hyperplasia at embryonic day 18.5, before exposure to microbes at birth (Lin et al., 2016; Newton et al., 2016). Based on our observations in *Ripk1*^EKO^ mice, we anticipate that the embryonic lethality of RIPK1 RHIM-mutant mice also requires nucleic acid sensing by ZBP1. It is not clear at which level RIPK1 operates to suppress ZBP1-mediated necroptosis. In the absence of an intact RHIM of RIPK1, RIPK3 spontaneously associates with ZBP1, suggesting that RIPK1 acts as a molecular scaffold to retain RIPK3 in an inactive state (Lin et al., 2016; Newton et al., 2016).

Others and we identified ZBP1 as a sensor of viral RNA (Maelfait et al., 2017; Sridharan et al., 2017; Thapa et al., 2016). Our current data brings forward the hypothesis that the detection of endogenous nucleic acids by ZBP1 contributes to inflammatory skin disease. Whether RNA or DNA serve as agonists for ZBP1 in RIPK1-deficient keratinocytes remains to be determined. Zα-domains found in ZBP1 and the RNA editing enzyme ADAR1 specifically bind to nucleic acids in the Z-conformation, including Z-RNA and Z-DNA (Herbert, 2019). How dsRNA or dsDNA are stabilised in the thermodynamically unfavourable Z-conformation in living cells is unknown. At least *in vitro*, alternating CG-rather than AT-sequences more readily transition into the Z-conformation suggesting that the sequence context is a contributing factor (Hall et al., 1984; Wang et al., 1979). Another possibility is that nucleic acid binding to ZBP1 promotes the stabilisation of the Z-conformer. Indeed, the second Zα2 domain of ZBP1 binds B-form DNA and facilitates its transition into Z-DNA (Kim et al., 2011a; Kim et al., 2011b). We cannot rule out that the mutation of the Zα-domains of ZBP1 affects other functions of ZBP1 apart from nucleic acid binding. However, our data showing that ectopically expressed Zα-domain mutant ZBP1 still activates NF-kB (Maelfait et al., 2017) and binds to RIPK3 (see Fig. S1C) suggests that at least the RHIM-RHIM interactions between ZBP1 and RIPK3 remained intact.

Complete *Ripk1* knockout mice die perinatally due to unconstrained activation of both apoptotic and necroptotic pathways (Dillon et al., 2014; Kaiser et al., 2014; Rickard et al., 2014b). At the epithelial barriers, there appears to be an interesting dichotomy in the survival functions of RIPK1. In intestinal epithelium, RIPK1 prevents cell death mediated by TNF and caspase-8-dependent apoptosis (Dannappel et al., 2014; Takahashi et al., 2014), whereas loss of RIPK1 in skin epithelial cells unleashes ZBP1-RIPK3-MLKL-mediated necroptosis (Dannappel et al., 2014). Why is one cell type more sensitive to apoptosis and another cell type more skewed towards necroptosis in the absence of RIPK1? A simple explanation may be the differential expression levels of components of cell death pathways such as FADD, caspase-8, RIPK3 and MLKL. However, this does not appear to be the case since keratinocyte-specific depletion of the linear ubiquitination chain assembly complex results in TNF/caspase-8-driven apoptosis driving dermatitis (Gerlach et al., 2011; Kumari et al., 2014; Rickard et al., 2014a; Taraborrelli et al., 2018). *Vice versa*, ablation of FADD or caspase-8 triggers necroptosis of intestinal epithelial cells leading to chronic intestinal inflammation (Gunther et al., 2011; Welz et al., 2011). A more complex scenario in which the mode of cell death is dictated by the availability of certain PRRs and their respective ligands together with regulatory mechanisms imprinted by the genetic background, the tissue environment and/or inflammatory cues is more likely.

In most cell types, including keratinocytes, ZBP1 levels are undetectable under steady state conditions and its transcription is strongly induced by IFNs (Shaw et al., 2017). Here, we demonstrate that the skin of *Ripk1*^EKO^ mice displays an innate antiviral immune response, which is responsible for the high levels of ZBP1 expression. Cultured RIPK1-deficient keratinocytes, however, do not express enhanced ZBP1 levels, indicating that keratinocyte-intrinsic triggers do not provide the cues stimulating the antiviral immune response in the skin of *Ripk1*^EKO^ mice. Bacterial or viral colonisation of the newborn epidermis during parturition may engage type-I IFN inducing nucleic acid sensors in keratinocytes or in adjacent immune cells, inducing ZBP1 levels above a certain toxic threshold. In addition, DAMPs released by necroptotic cells can be detected by PRRs in neighbouring cells, further supporting inflammation. Once the initial trigger is delivered, RIPK1-deficient keratinocytes undergoing ZBP1-dependent necroptosis may set in motion an auto-amplifying inflammatory signalling cascade establishing chronic inflammation. This is reminiscent of the chronic proliferative dermatitis phenotype of SHARPIN-deficient mice, where the initial trigger is TNF-induced keratinocyte cell death. Release of dsRNA from dying cells then activates TLR3, which in SHARPIN-deficient conditions causes cell death resulting in a self-promoting cycle of inflammation (Zinngrebe et al., 2016).

We demonstrate that keratinocyte necroptosis drives IL-17-mediated inflammation. A recent study showed that the transgenic expression of cFLIPS, a splice variant of the cFLIPL gene *Cflar*, caused the activation of ILC3s, which depended on necroptosis of intestinal epithelial cells (Shindo et al., 2019). At this stage, we do not know whether the IL-17 response is protective and aiding in tissue repair or whether it contributes to disease progression and chronic skin inflammation, as seen in patients with psoriasis (Brembilla et al., 2018; Ho and Kupper, 2019). Future studies in IL-23 or IL-17 deficient animals will provide clarity in this matter (Tait Wojno et al., 2019). Finally, our observations may assign important roles for endogenous nucleic acid sensing by ZBP1 in primary immunodeficiency and inflammatory disease observed in RIPK1 deficient patients (Cuchet-Lourenco et al., 2018; Li et al., 2019).

## Materials and methods

### Mice

*Ripk1*^FL/FL^ (Takahashi et al., 2014), Keratin-5-Cre (Ramirez et al., 2004), *Ripk3*^−/−^ (Newton et al., 2004), *Mlkl*^FL/FL^ (Murphy et al., 2013), *Zbp1*^−/−^ (Ishii et al., 2008), *Casp1/11*^−/−^ (Kuida et al., 1995) and *Zbp1*^Zα1α2/Zα1α2^ (Maelfait et al., 2017) mice were housed in individually ventilated cages at the VIB-UGent Center for Inflammation Research in a specific pathogen-free facility. *Casp1/11*^−/−^ mice were purchased from The Jackson Laboratory (B6N.129S2-Casp1^tm1Flv^/J). *Ripk1*^FL/FL^, *Mlkl*^FL/FL^ and *Zbp1*^Zα1α2/Zα1α2^ mice were generated in C57Bl/6 ES cells, Keratin-5-Cre transgenes were generated in C57Bl/6JxDBA/2J ES cells and *Ripk3*^−/−^, *Casp1/11*^−/−^ (Kayagaki et al., 2015) and *Zbp1*^−/−^ (Koehler et al., 2020) mice were generated in 129-derived ES cells. All lines were maintained in a C57Bl/6 background. Mouse lines that were not generated in C57Bl/6 ES cells were backcrossed at least 10 times to a C57Bl/6 background. Littermates were used as controls for all experiments. All experiments were conducted following approval by the local Ethics Committee of Ghent University.

### Reagents

Mouse TNF and mouse IFNγ were produced by the VIB Protein Service Facility. zVAD-fmk (Bachem; BACEN-1510.0005), CHX (Sigma; C7698), poly(I:C) (Invivogen; tlrl-pic), mouse IFNβ (PBL Biomedical Laboratories) and mouse CXCL10 ELISA (ThermoFisher; BMS6018MST) were obtained from commercial sources.

### Western blotting and immunoprecipitation

For Western blotting, cells were washed with PBS and lysed in protein lysis buffer (50 mM Tris.HCl pH 7.5, 1% Igepal CA-630, 150 mM NaCl) supplemented with complete protease inhibitor cocktail (Roche, 11697498001) and PhosSTOP (Roche, 4906845001). Lysates were cleared by centrifugation at 16,000 g for 15 min and 5X Laemlli loading buffer (250 mM Tris.HCl pH 6.8, 10% SDS, 0,5% Bromophenol blue, 50% glycerol, 20% β-mercaptoethanol) was added to the supernatant. Finally, samples were incubated at 95°C for 5 min and analysed using Tris-Glycine SDS-PAGE and semi-dry immunoblotting. For immunoprecipitation, HEK293T cells were transfected with Lipofectamine 2000 (Life Technologies) with N-terminally V5-tagged mouse ZBP1 and mouse RIPK3 cloned into pcDNA3 expression vectors. 24 hours post-transfection, cells were washed in PBS and lysed in protein lysis buffer. Lysates were cleared from debris by centrifugation at 16,000 g for 15 min and 10% of the sample was used for input control. The remaining lysate was incubated overnight at 4 °C on a rotating wheel using anti-V5 Affinity Gel beads (Sigma, A7345). Beads were washed three times with protein lysis buffer. Finally, the beads were resuspended in 1X Laemmli buffer and incubated at 95°C for 5 min Primary antibodies used in this study are anti-ZBP1 (Adipogen, Zippy-1), anti-RIPK1 (CST, 3493), anti-RIPK3 (ProSci, 2283), anti-MLKL (Millipore, MABC604), anti-phospho-Thr231/Ser232-RIPK3 (CST, 57220), anti-phospho-S345-MLKL (Abcam, 196436), anti-V5-HRP (Invitrogen, R960-25) and anti-β-Tubulin-HRP (Abcam, 21058).

### Histology and immunohistochemistry

Back skin biopsy specimens were fixed in 4% paraformaldehyde in phosphate buffered saline overnight at 4 °C, after which they were embedded in paraffin and sectioned at 5 μm thickness. Sections were deparaffinised before hematoxylin and eosin (H&E) staining using a Varistain Slide Stainer. For determining the epidermal thickness, H&E stained skin sections with a length of approximately 1 cm were imaged with a ZEISS Axio Scan slide scanner. Then, the thickness of the epidermis was measured at 10 points per skin section with the ZEISS Blue software and the values were expressed as the average of these 10 measurements. For immunofluorescence, sections were deparaffinised and rehydrated using a Varistain Slide Stainer. Antigen retrieval was performed by boiling sections at 95°C for 10 min in antigen retrieval solution (Vector, H-3301). Slides were then treated with 3% H_2_O_2_ in PBS for 10 min and 0.1 M NaBH_4_ in PBS for 2 hours at room temperature to reduce background. After washing in PBS, tissues were blocked in 1% BSA and 1% goat serum in PBS for 30 min at room temperature. Tissue sections were stained overnight at 4°C with following primary antibodies: anti-K6 (Covance, PRB-169P), anti-K5 (Covance, PRB-160P), anti-K1 (Covance, PRB-165P) and anti-ZBP1 (Adipogen, Zippy-1). Next, sections were incubated with donkey anti-mouse CF633 (Gentaur, 20124-1) or goat anti-rabbit Dylight488 (ThermoFisher; 35552) and DAPI (Life Technologies, D1306) for 30 min at room temperature. Images were acquired on a Zeiss LSM880 Fast AiryScan confocal microscope using ZEN Software (Zeiss) and processed using Fiji (ImageJ).

### RT-qPCR

Snap-frozen back skin was homogenised on dry ice using a mortar and pestle. Total RNA was purified using RNeasy columns (Qiagen) with on-column DNase I digestion. cDNA synthesis was performed using the SensiFast cDNA synthesis kit (Bioline, BIO-65054). SensiFast SYBR No-Rox kit (Bioline, BIO-98005) or PrimeTime qPCR Master Mix (IDT, 1055771) were used for cDNA amplification using a Lightcycler 480 system (Roche). The following primers were used for SYBR-green based detection and the median expression of *Rpl13a* and *Hprt* were used for normalisation: *Ifng* forward 5’GCCAAGCGGCTGACTGA and reverse 5’TCAGTGAAGTAAAGGTACAAGCTACAATCT; *S100a8* forward 5’GGAGTTCCTTGCGATGGTGAT and reverse 5’CAGCCCTAGGCCAGAAGCT; *Tnf*, forward 5’CCACCACGCTCTTCTGTCTA and reverse 5’GCTACAGGCTTGTCACTCGAA; *Il33*, forward 5’GAGCATCCAAGGAACTTCAC and reverse 5’AGATGTCTGTGTCTTTGA; *Isg15*, forward 5’TGACGCAGACTGTAGACACG and reverse 5’TGGGGCTTTAGGCCATACTC;*Ifit1*,forward 5’CAGAAGCACACATTGAAGAA and reverse 5’TGTAAGTAGCCAGAGGAAGG; *Ccl20*,forward 5’TGCTATCATCTTTCACACGA and reverse 5’CATCTTCTTGACTCTTAGGCTG; *Il6*, forward 5’TTCTCTGGGAAATCGTGGAAA and reverse 5’TCAGAATTGCCATTGCACAAC;*Il1a*,forward 5’CCTGCAGTCCATAACCCATGA and reverse 5’ACTTCTGCCTGACGAGCTTCA; *Il1b*, forward 5’CCAAAAGATGAAGGGCTGCTT and reverse 5’TCATCAGGACAGCCCAGGTC; *Rpl13a*, forward 5’CCTGCTGCTCTCAAGGTTGTT and reverse 5’TGGTTGTCACTGCCTGGTACTT and *Hprt*, forward 5’CAAGCTTGCTGGTGAAAAGGA and reverse 5’TGCGCTCATCTTAGGCTTTGTA. The following oligo’s were purchased from IDT for probe-based detection and the median expression of *Tbp* and *Actb* were used for normalisation: *Zbp1*, Mm.PT.58.21951435; *Ifi44*, Mm.PT.58.12162024, *Tbp*, Mm.PT.39a.22214839 and *Actb*, Mm.PT.39a.22214843.g.

### Primary keratinocytes cultures

Primary keratinocytes were isolated from back skin of 4 to 5 weeks old mice. Back skin was shaved with electrical clippers and sterilized with 10% betadine and 70% ethanol. Subcutaneous fat and muscles were removed by mechanical scrapping. Subsequently, the skin was carefully placed on 0.25% trypsin (Gibco, 25050014) with dermal side downwards for 2 hours at 37°C. Epidermis was separated from the dermis with a forceps, cut into fine pieces, and incubated in fresh 0.25% trypsin for 5 minutes at 37°C with rotation. After neutralization with supplemented FAD medium (see below), the epidermal cell suspension was filtered through a 70 µm cell strainer. The keratinocytes were seeded on J2 3T3 feeder cells that were mitotically inactivated by treatment with 4 µg/ml Mitomycin C (Sigma-Aldrich, #M-0503) for two hours, in 1 µg/ml collagen I-coated falcons (Sigma-Aldrich, 5006). Keratinocytes were cultured at 32°C in a humidified atmosphere of 5% CO_2_ and medium was replaced every two days. After 12-14 days the remaining J2 3T3 feeder cells were removed from the culture by short incubation with 0.25% trypsin and the keratinocytes were harvested subsequently. Keratinocytes were seeded at a density of 25,000 cells per cm^2^ into collagen I-coated culture plates for cell death measurements (48-well) and Western blot analysis (12-well). Primary mouse keratinocytes were cultured in custom-made (Biochrom, Merck) DMEM/HAM’s F12 (FAD) medium with low Ca^2+^ (50 µM) supplemented with 10% FCS (Gibco) treated with chelex 100 resin (Bio-Rad, 142-2832), 0.18 mM adenine (Sigma-Aldrich, A2786), 0.5 µg/ml hydrocortisone (Sigma-Aldrich, H4001), 5 µg/ml insulin (Sigma-Aldrich, I3536), 10^−10^ M cholera toxin (Sigma-Aldrich, C8052), 10 ng/mL EGF (ThermoFisher Scientific, 53003018), 2 mM glutamine (Lonza, BE17-605F), 1 mM pyruvate (Sigma-Aldrich, S8636), 100 U/ml penicillin and 100 µg/ml streptomycin (Sigma-Aldrich, P4333), and 16 µg/ml gentamycin (Gibco, 15710064).

### Cell death assays

25,000 primary keratinocytes were seeded per well in 1 µg/ml collagen I-coated 48-well plates in FAD medium. 24 hour later, the cell impermeable dye SYTOX Green (1 µM, ThermoFisher Scientific, S7020) was added to the culture medium together with the indicated stimuli. SYTOX Green uptake was imaged every 2 or 3 hour with an IncuCyte Live-Cell Analysis system (Essen BioScience) at 37°C. The relative percentage of SYTOX Green cells was determined by dividing the number of SYTOX Green positive cells per image by the percentage of confluency (using phase contrast images) at every time point.

### Keratinocyte transduction

HEK293T were cultured in high-glucose (4500 mg/L) DMEM (Gibco, 41965-039) supplemented with 10% FCS and 2 mM glutamine (Lonza, BE17-605F). For lentivirus production, HEK293T cells were transfected with N-terminally 3XFLAG and V5-tagged wild type mouse ZBP1 or Zα1α2-mutant mouse ZBP1 transducing vectors in the pLenti6 backbone (Life Technologies) (Maelfait et al., 2017) together with the pCMV delta R8.91 gag-pol expressing packaging plasmids and pMD2.G VSV-G expressing envelope plasmid. 24 hour post-transfection, cells were washed and FAD medium was added. 48 hour post-transfection, the viral supernatant was harvested and used to transduce 25,000 mouse primary keratinocytes seeded in 48-well plates in the presence of 8 µg/ml polybrene (Sigma Aldrich). The next day, viral particle-containing medium was removed and cell death was measured for 48 hours as described above.

### Skin processing for flow cytometry analysis

A piece of shaved mouse skin (± 12 cm^2^) was isolated from 4-5 weeks old mice. Subcutaneous fat and muscles were removed by mechanical scrapping with a scalpel. Subsequently, the skin was carefully placed on 0.4 mg/ml Dispase II (Roche, 4492078001) with the dermal side facing downwards for 2 hours at 37°C. Epidermis was separated from the dermis with a forceps, cut into fine pieces, and incubated in 2 ml enzymatic solution containing 1.5 mg/ml collagenase type IV (Worthington, LS004188) and 0.5 mg/ml Dnase I (Roche, 10104159001) for 20 minutes at 37°C with shaking. After neutralization with 2% FCS RMPI medium, the cell suspension was filtered through a 70 µm cell strainer to obtain a single cell suspension. For detection of intercellular cytokines, *in vitro* activation was performed. Single cells suspension from epidermis was seeded into 96-well U-bottom plate in RPMI 1640 medium supplemented with 10% FCS, 2 mM glutamine (Lonza, BE17-605F), 1 mM sodium pyruvate (Sigma-Aldrich, S8636), 100 U/ml penicillin and 100 µg/ml streptomycin (Sigma-Aldrich, P4333), 50 mM β-mercaptoethanol (Gibco, 31350-010). Subsequently, the cell suspension was stimulated with eBioscience Cell Stimulation Cocktail plus protein transport inhibitor (eBioscience, 00-4975-03) for 4 hours at 37°C in a humidified 5% CO2 incubator.

### Flow cytometry

Single cell suspensions were first stained with anti-mouse CD16/CD32 (Fc-block; BD Biosciences; 553142) and dead cells were excluded with the Fixable Viability Dye eFluor506 (eBioscience; 65-0866-14) for 30 min at 4°C in PBS. Next, cell surface markers were stained for 30 min at 4°C in FACS buffer (PBS, 5% FCS, 1mM EDTA and 0.05 sodium azide). For intracellular cytokine analysis, cells were fixed for 20 minutes at 4°C using BD Cytofix/Cytoperm (BD Biosciences, 554714) and washed twice with BD Perm/Wash buffer (BD Biosciences, 554714). Intracellular IFNγ and IL-17A were stained in BD Perm/Wash Buffer for 30 min at 4°C. Cells were acquired with on an LSR Fortessa or a FACSymphony (BD Biosciences) and data were analysed with FlowJo software (TriStar). The total number of cells were counted using a FACSVerse (BD Biosciences). The following fluorochrome-conjugated antibodies were used: CD3e#BUV395 (1/100; 563565), CD161#BV605 (1/300; 563220), CD4#FITC (1/400; 557307), TCRγδ#PECF594 (1/500; 563532), CD11b#PerCP-Cy5.5 (1/1000; 550993), and CD44#PE (1/400; 553134) were from BD Biosciences. CD8α#eFluor450 (1/400; 48-0081-82), CD45#AF700 (1/200; 56-0451), MHC Class II (I-A/I-E)#FITC (1/1000; 11-5321), CD11c#PE-Cy7 (1/500; 25-0114-82) and IL17A#APC (1/200; 17-7177-81) were from eBioscience. CD19#BV785 (1/400; 115543), CD11c#BV785 (1/200; 117336), Ter-119#BV785 (1/400; 116245), Ly-6G#BV785 (1/400; 127645), TCRβ#APC-Cy7 (1/200; 109220), CD64(FcγRI)#BV711 (1/100; 139311) and IFNγ#PE/Cy7 (1/300; 505825) were from Biolegend.

### Statistical analyses

Statistical analyses were performed using Prism 8.2.1 (GraphPad Software). Statistical methods are described in the figure legends.

## Online supplemental material

Figure S1 provides additional information for figure 1 and shows pictures of skin lesions of *Ripk1*^EKO^ *Zbp1*^+/+^ and *Ripk1*^EKO^ *Zbp1*^Zα1α2/Zα1α2^ mice, demonstrates equal protein expression of ZBP1 and Zα-domain mutant ZBP1 in RIPK1-deficient keratinocytes, and shows lesion formation of *Ripk1*^EKO^ mice crossed to *Zbp1*^−/−^ and *Ripk3*^−/−^ animals. Figure S2 relates to figure 2 and figure 3 and shows the gating strategy for figure 2A and additional RT-qPCR analyses of ISGs and pro-inflammatory genes in the skin of *Ripk1*^EKO^ *Zbp1*^+/+^ and *Ripk1*^EKO^ *Zbp1*^Zα1α2/Zα1α2^ mice. Figure S3 relates to figure 4 and describes cell death induction of RIPK1-deficient primary keratinocytes after transduction with ZBP1 or after treatment with TNF + zVAD or TNF + CHX. Figure S3 also shows that ISG expression in keratinocytes upon IFNγ treatment is not affected by RIPK1 deficiency or Zα-domain mutation of ZBP1.

## Author contributions

M. Devos, G. Tanghe, P. Vandenabeele, W. Declercq and J. Maelfait designed the study. M. Devos, G. Tanghe, B. Gilbert, E. Dierick, M. Verheirstraeten, J. Nemegeer, R. de Reuver, S. Lefebvre, J. De Munck, and J. Maelfait carried out the experiments. J. Rehwinkel provided critical reagents and scientific advice. M. Devos, G. Tanghe, B. Gilbert, W. Declercq and J. Maelfait analysed the results. M. Devos, G. Tanghe, W. Declercq and J. Maelfait wrote the manuscript.

## Acknowledgements

We are grateful to Kim Newton and Vishva Dixit for providing *Ripk3*^−/−^ mice, to James Murphy and Warren Alexander for providing *Mlkl*^FL/FL^ mice, and to Ken Ishii and Shizuo Akira for providing *Zbp1*^*-/-*^ animals. We thank the members of the VIB Flow Core, Protein Service Facility and the Microscopy Core, and Kelly Lemeire for technical assistance. We thank members of the Rehwinkel and Vandenabeele lab, Sophie Janssens and Mathieu Bertrand for helpful discussions. This research would not have been possible without support from the following funding agencies. J. Maelfait was supported by and Odysseus II Grant (G0H8618N) from the Research Foundation Flanders and by Ghent University. The W. Declercq lab was supported by ‘Vlaams Instituut voor Biotechnologie’ (VIB), a UGent Grant (GOA-01G01914) and Stichting tegen Kanker (FAF-F/2016/868). Research in the P. Vandenabeele unit is supported by Belgian grants (EOS 30826052 MODEL-IDI), Flemish grants (FWO G.0C31.14N, G.0C37.14N, FWO G0E04.16N, G.0C76.18N, G.0B71.18N, G0B9620N), grants from Ghent University (BOF16/MET_V/007 Methusalem grant), Foundation against Cancer (FAF-F/2016/865) and VIB. J. Rehwinkel is funded by the UK Medical Research Council (MRC core funding of the MRC Human Immunology Unit). The authors declare no competing financial interest.

---

**Supplemental Figure S1.**
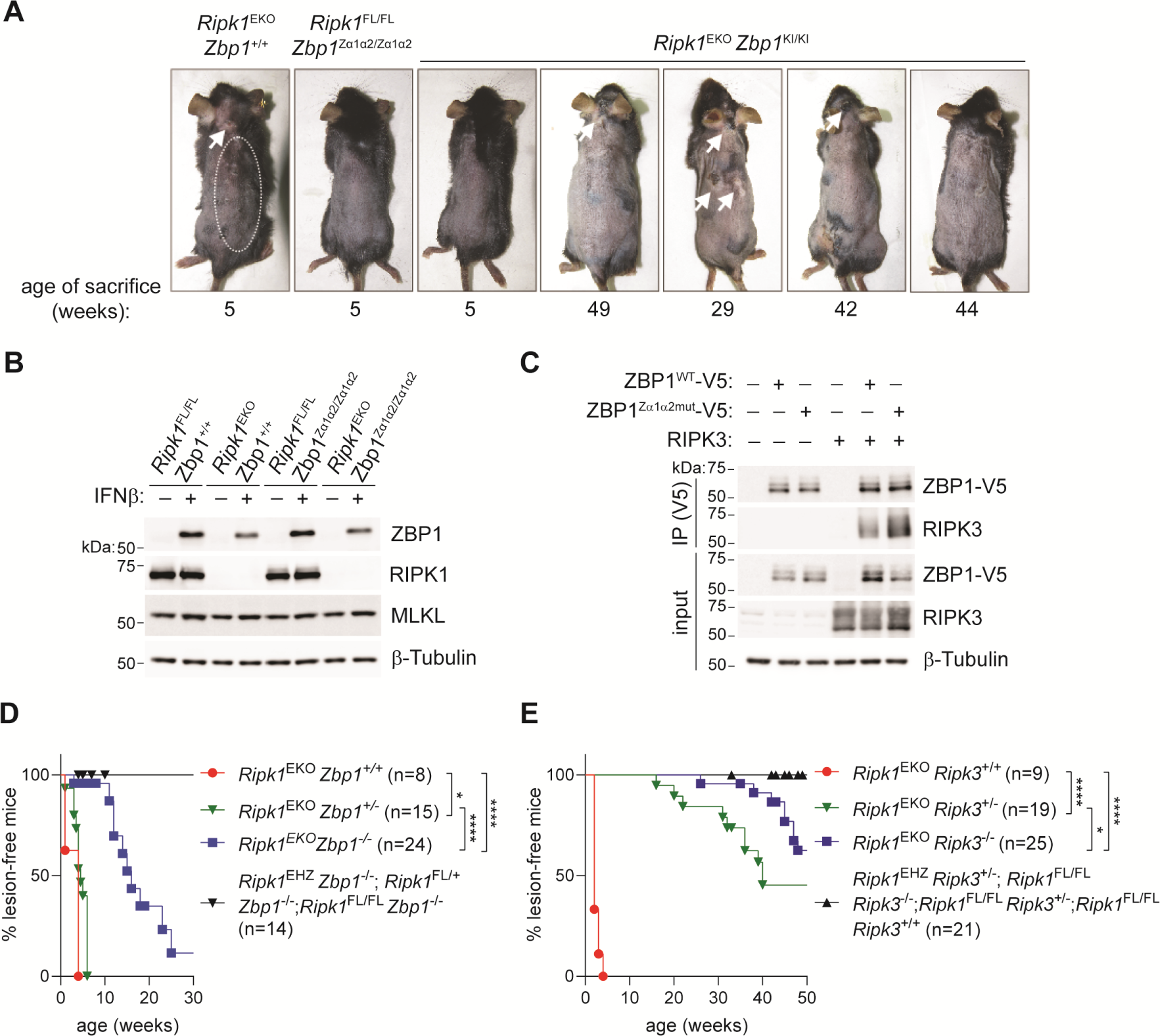
Skin pathology of *Ripk1*^EKO^ mice is dependent ZBP1 and RIPK3. **(A)** Macroscopic appearance of shaven mice of the indicated genotypes at the indicated age. Lesional skin is depicted by white arrows or by the dotted line. **(B)** Protein expression analysis by Western blotting on lysates of primary keratinocytes isolated from mice of the indicated genotypes. Cells were stimulated for 24 hours with 200 U/ml IFNβ to induce ZBP1 expression. **(C)** N-terminally V5-tagged wild type mouse ZBP1 (ZBP1^WT^-V5) or Zα-domain mutant ZBP1 (ZBP1^Zα1α2mut^-V5) and mouse RIPK3 were expressed in HEK293T cells. 24 hours later, the presence of ZBP1 and RIPK3 in V5-immunoprecipitates (IP) were analysed by Western blotting. Protein expression in input samples is shown in the lower panels. **(D-E)** Kaplan-Meier plots of lesion appearance of *Ripk1*^EKO^ mice crossed to (D) ZBP1-deficient (*Zbp1*^−/−^) or (E) RIPK3-deficient (*Ripk3*^−/−^) animals. *p < 0.05, ****p < 0.0001 by Log-Rank test. Data shown (B) and (C) are representative of at least 2 independent experiments.

**Supplemental Figure S2.**
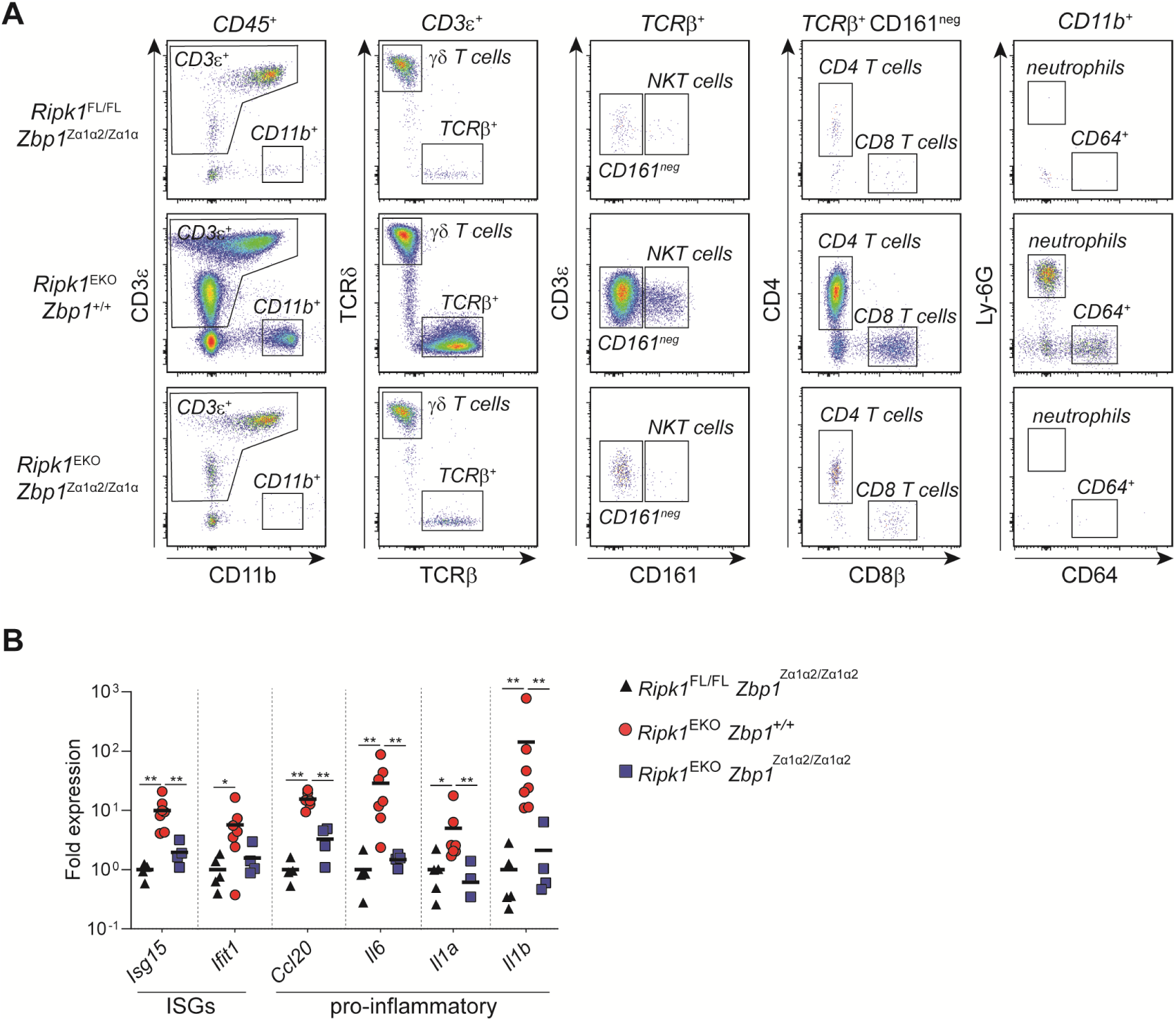
ZBP1 activation drives antiviral and IL-17 immune responses in *Ripk1*^EKO^ mice. **(A)** Flow cytometry gating strategy of CD45^+^ immune cell subsets isolated from mice of the indicated genotypes and quantified in figure 2A. **(B)** RT-qPCR analysis of the indicated interferon-stimulated genes (ISGs) and pro-inflammatory genes in whole back skin of 4 to 5 week old mice of the indicated genotypes. Data are representative of 2 independent experiments. *p < 0.05, **p < 0.01 by Mann-Whitney U-test.

**Supplemental Figure S3.**
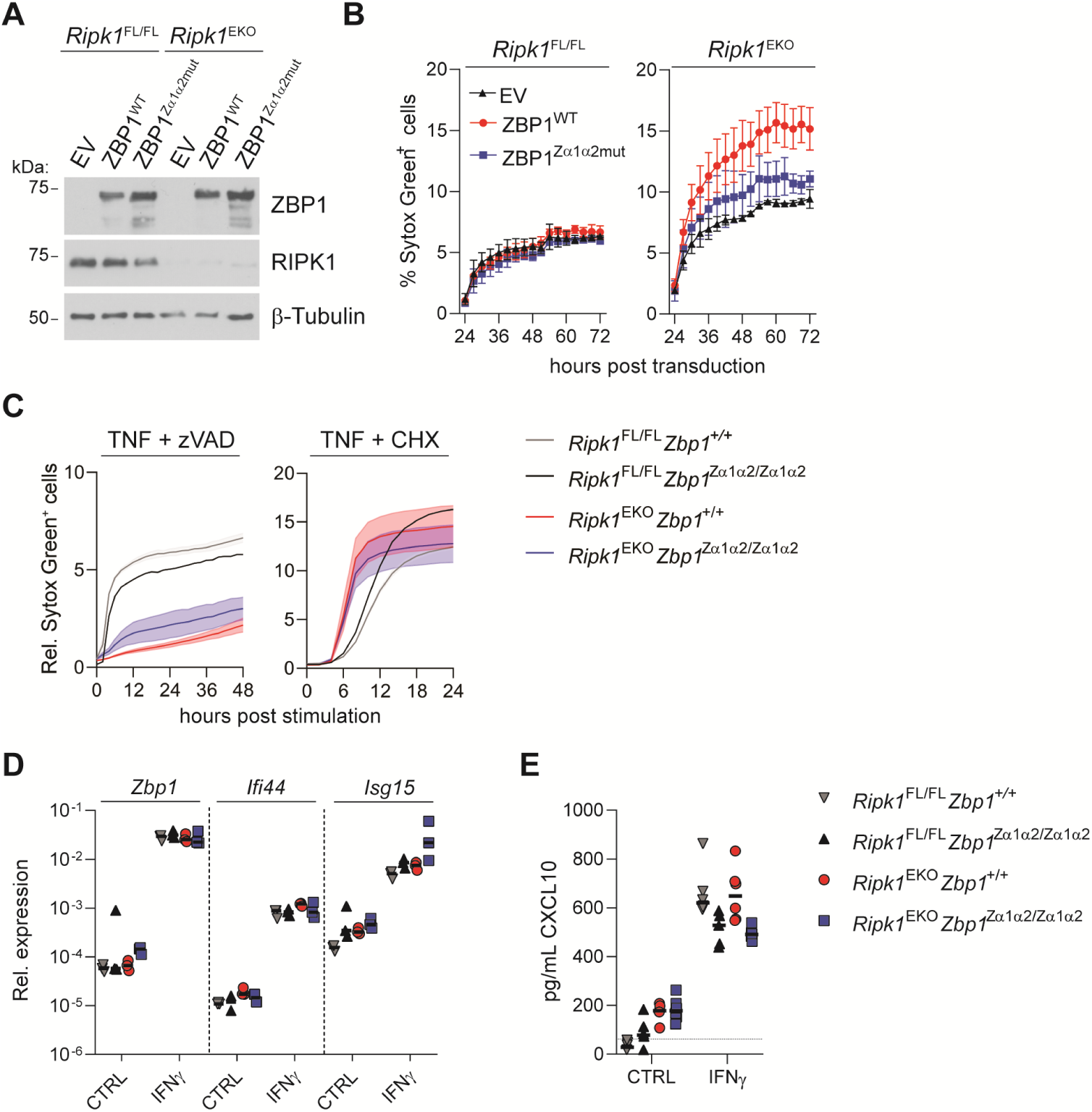
ZBP1 activation by endogenous nucleic acids induces necroptosis in RIPK1-deficient keratinocytes. **(A)**Primary keratinocytes isolated from the indicated genotypes were transduced with empty vector (EV), wild type mouse ZBP1 (ZBP1^WT^) or ZBP1 in which the two Zα-domains were mutated (ZBP1^Zα1α2mut^). 48 hours post-transduction, protein expression of ZBP1 and RIPK1 was analysed by Western blotting. ZBP1 was N-terminally fused to a 3XFLAG and V5-tag. **(B)** Analysis of cell death by measuring Sytox Green uptake on primary keratinocytes, transduced as described in (B). Measurements were performed every 3 hours starting at 24 hour until 72 hours post-transduction. **(C)** Analysis of cell death upon stimulation with 30 ng/ml TNF with 20 µM zVAD-fmk (TNF + zVAD) or 30 ng/ml TNF with 5 µg/ml cycloheximide (TNF + CHX) by measuring Sytox Green uptake every 2 hours of primary keratinocytes isolated from mice of the indicated genotype. Solid lines represent the mean of cell death curves of 1 to 4 primary keratinocyte cultures isolated from different mice. Shaded areas indicate SEM. **(D)** RT-qPCR analysis of the indicated interferon-stimulated genes (ISGs) after 18 hours stimulation with 200 U/ml IFNγ of primary keratinocytes isolated from mice of the indicated genotypes. **(E)** Measurement of CXCL10 protein levels in supernatant collected from cells from (D) by ELISA. The dotted line indicates the detection limit. Data in (A), (B), (C), (D) and (E) are representative of 2 independent experiments.

